# Culture Positivity Reflects Quantitative Bacterial Bioburden in Open Fractures

**DOI:** 10.64898/2026.09.22.753238

**Authors:** Emily Ann McClure, Roman Natoli, Renan C. Castillo, Anthony R. Carlini, Joseph Wenke, Robert V. O’Toole, Gregory M. Schrank, William Obremskey, Stephen Warner, Michael Bosse, Benjamin D. Ross, Leah Gitajn, METRC

## Abstract

**Background:** Infection following high-energy open fractures remains a major challenge. Open fractures are exposed to environmental and skin pathogens, which, along with traumatized tissue and implanted hardware, create a favorable environment for infection. 24-83% of open fractures yield positive culture results. Despite culture-positive wounds being at higher risk for infection, culture data are rarely used in clinical decision-making. Culture-positivity in open fractures is believed to indicate higher wound bioburden, but this hypothesis remains unconfirmed. This study aimed to correlate culture positivity and quantitative wound bioburden using next-generation sequencing (NGS) and to characterize patterns of bioburden and bacterial diversity amongst patients with open fracture and Fracture Related Infection. We hypothesized that higher wound bioburden would be associated with culture positivity.

**Methods:** This secondary analysis of the *METRC Bioburden Study* included 78 patients with severe open tibia fractures who underwent standardized tissue sampling at definitive wound closure and at later fracture site revision surgery. DNA extracted from intraoperative tissue was analyzed via 16S rRNA gene amplicon sequencing. Quantitative bioburden metrics included bacterial read percentage and bacteria-to-human read ratio (BHR). Associations between sequencing-based bioburden, culture results, and clinical infection (CDC-defined) were evaluated using Mann– Whitney U tests, McNemar’s chi-squared tests, and unsupervised clustering analyses.

**Results:** Culture-positive samples exhibited significantly higher bacterial read percentages (17.5 ± 26.7% vs. 0.8 ± 4.7%, *p* = 8×10□□) and higher log(BHR) values (1.73 ± 10.35 vs. 0.06 ± 0.52, *p* = 3×10□□) than culture-negative samples. Optimal thresholds yielded 65% sensitivity/94% specificity (bacterial reads) and 89% sensitivity/69% specificity (BHR) for predicting culture positivity. Neither baseline culture nor NGS positivity predicted later infection. However, follow-up samples from patients with established infection showed significantly higher bioburden and distinct microbiome structures. Clustering analyses identified two infection-associated microbial profiles: one with high bioburden and high diversity (polymicrobial infection) and one with low bioburden and low diversity (single-dominant pathogen).

**Conclusions:** Culture positivity in open fractures correlates strongly with increased bacterial bioburden, indicating that positive cultures may primarily reflect microbial load rather than specific pathogens. Quantitative NGS-based measures, including bacterial read proportion and BHR, may offer a biologically grounded approach to assess wound contamination and may inform infection risk. Distinct bioburden–diversity patterns suggest heterogeneous infection phenotypes that may require tailored surgical and antimicrobial strategies.

## INTRODUCTION

Infection following high-energy open fracture remains a major challenge for both patients and orthopaedic surgeons.^1–8^ Fracture related infections carry an estimated annual economic burden of $1.8 billion in the United States alone.^9–13^ Open fractures, which are exposed to the external environment, often harbor dirt and pathogens from the patient’s skin, the environment, and hospital-acquired bacteria. The combination of devitalized bone and soft tissue and the implantation of hardware creates conditions that favor pathogen persistence and biofilm formation, contributing to the high rates of infection in this patient population.^14^

Culture positivity at time of debridement or closure is common, and reported to occur in 24 to 83% of open fractures. ^15–19^ Culture positivity at definitive wound closure is clinically relevant, as it is associated with increased risk for subsequent infection.^15,17,18^ However, the pathogens responsible for later infection often differ from those identified initially,^15,17,18^ making the early culture results difficult to interpret and limiting their clinical utility. Although culture positivity has been proposed as a tool to guide repeat debridement and prophylactic antibiotic choice,^19–21^ this strategy remains inconsistently applied and few surgeons routinely obtain culture during the initial debridement or definitive closure.^15,17,19,22,23^

One proposed explanation for the inconsistency between initial culture positivity and infection outcome is that culture positivity may simply reflect a higher overall wound bioburden compared to culture negative cases.^19^ However, this hypothesis has not been confirmed. In this study, we aimed to evaluate whether bacterial bioburden, quantified using 16S rRNA gene amplicon sequencing, correlates with culture positivity and subsequent infection in open fractures. This approach offers the potential for a more sensitive and comprehensive alternative to traditional culture and may help clarify this relationship. We hypothesized that higher bioburden would be associated with both culture positivity.

## MATERIALS AND METHODS

### Study design

This is a secondary analysis of the Bioburden Study (NCT01496014).^20^ Bioburden was a multicenter, prospective cohort study of the terminal wound culture results in severe open tibia fractures that were treated with delayed wound closure or coverage. Patients aged 18 and 74 were eligible for inclusion if they sustained either a Gustilo type III tibial fracture (plateau, shaft or pilon) requiring repeat debridement following initial debridement and stabilization, or a traumatic transtibial amputation requiring delayed primary closure or flap coverage. At the time of definitive wound closure (baseline), all patients underwent standardized specimen collection, including swab samples obtained for routine microbiologic culture and biobanking for future analyses. Additional samples were collected if corrective surgery was performed (follow-up). The clinical microbiology and tissue sample collection protocols are summarized in Appendix 1 of the Bioburden paper.^20^ All patients were treated according to the standard of care of each participating center, without assigned specific treatment interventions. Patients were prospectively monitored for 12 months. All outcomes, including Surgical Site Infection using the Centers for Disease Control and Prevention (CDC) definition,^24^ were adjudicated. A total of 646 patients were enrolled into the Bioburden Study including 459 patients with definitive wound closure culture data.

Patients for this study were randomly selected from those with the most frozen tissue aliquots. Efforts were made to balance number of randomly selected samples to include three groups 1) culture positive infections, n=40; 2) culture negative infections, n=8; and 3) uninfected patients, n=43. Random samples were used as sequencing controls between sequencing runs.

Complete patient demographic information and wound characteristics are listed in Tables 1 and 2 respectively.

**Table 1.** Qualitative measure of bioburden. Values represent count of positive samples grouped by timepoint, infection diagnosis, NGS result, and culture result. (NGS=16S rRNA gene v1-3 amplicon sequencing). The relationship between culture and NGS-positivity was significant at both baseline and follow-up (McNemar’s Chi-squared test, p=0.043 and p=0.009 respectively).

|  |  | Baseline |  |  | Follow-up |  |  |
| --- | --- | --- | --- | --- | --- | --- | --- |
|  |  | NGS negative | NGS positive | Total | NGS negative | NGS positive | Total |
| UNINFECTED | Culture negative | 6 | 2 | 8 | 2 | 4 | 6 |
|  | Culture positive | 0 | 15 | 15 | 0 | 14 | 14 |
|  | Total | 6 | 17 | 23 | 2 | 18 | 20 |
| INFECTION | Culture negative | 5 | 2 | 7 | 0 | 1 | 1 |
|  | Culture positive | 4 | 4 | 8 | 2 | 30 | 32 |
|  | Total | 9 | 6 | 15 | 2 | 31 | 33 |

**Table 2.** Number of patients diagnosed with an infection predicted by alpha metrics, bioburden, or both. In total, 27/35 infections were correctly identified and only 2/17 uninfected samples were mis-identified as infected.

|  | Alpha prediction |  | logBHR prediction | Dual prediction |  | No prediction |
| --- | --- | --- | --- | --- | --- | --- |
|  | low | high | high | low | high |  |
| infection | 9 | 12 | 12 | 9 | 18 | 8 |
| uninfected | 2 | 0 | 0 | 2 | 0 | 17 |

### DNA extraction, sequencing, and analysis

Tissues were removed from −80°C storage, thawed, and RNA later removed by aspiration. Tissue was homogenized using a Seward Stomacher. 25-100mg of homogenized tissue was then transferred to a microcentrifuge tube and DNA was extracted from 25-100mg of homogenized tissue using the Qiagen DNeasy Blood and Tissue Kit with modifications for the complete lysis of tissue. Samples with added 270µL buffer ATL and 30µL proteinase K were incubated at 56°C for 12-36h for complete lysis. Incubated samples were then combined with 50µL 0.1mm Zirconia beads and 50µL 0.7mm Zirconia beads and homogenized for 10min at 25Hz using the Qiagen Tissuelyser. Samples were then centrifuged at 10,000rpm for 1min. 600µL of ethanol:buffer AL solution was then added and entire mixture was vortexed and transferred to a DNeasy spin column. Protocol continued as listed for the DNeasy Blood and Tissue Kit. Samples eluted into buffer AE were stored at −20°C. Purified DNA was amplified via PCR using primers flanking the V1-V3 region of the 16S rRNA gene. Amplified DNA was sequenced on an Illumina MiSeq. Resulting community sequences were processed using R (version 4.5.0). Initial ASV assignments were processed using the standard DADA2 pipeline.^25,26^ Primers were trimmed then sequences filtered: 0 Ns, truncated quality score <2, and maximum expected error rate of 3. Forward and reverse sequences were merged and chimeras removed. ASVs were assigned taxonomy by comparing to the SILVA SSU 132 data set.^27^ Any ASVs not identified to the genus level and considered significant in later analysis were compared to the BLAST nucleotide data set for taxonomic assignment.

The untrimmed 16S rRNA gene amplicon dataset consists of 34 negative controls and 92 unique samples. Amplicon size was limited to 415 to 505bp to reach a data set containing 3,514 (792 bacterial) unique ASVs. Negative controls were used to identify and trim contaminant ASVs from the dataset using the decontam package. Rarefaction curves were prepared for each sample individually to determine minimum sampling depth of 350 reads. Any samples with fewer than 350 bacterial reads were considered “failed” (having 0 reads). The resulting dataset contained 745 bacterial ASVs and 72 samples. Samples contained median 11,572 (max 191,353) bacterial reads per sample: Individual bacterial ASVs were present at median 197 (max 146,202) reads (Figure S1).

### Bioburden calculations

Since primers targeting the V1-V3 region of the bacterial 16S rRNA gene can amplify off-target sequences in the human genome, ASVs assigned to order Primates were used to estimate the bacteria-to-human ratio (BHR). BHR is one indirect measure of wound bioburden, with a low proportion of human-derived reads indicating high bacterial bioburden.^28^ The abundance of the most prevalent human ASVs (prevalence in the 99.5% quartile) were then compared to the BLASTn database to confirm they were *Homo sapiens*. The most prevalent human ASVs were then used as a proxy for all human ASVs. Human ASVs ranged from 250 to 510bp (72 ASVs).

### Statistical Analysis

McNemar’s chi-squared test was used to compare the relationship between qualitative outcomes. Youden’s J statistic was used to calculate optimum thresholds to separate groups using the cutpointr package (version 1.2.1).^34^ A Mann-Whitney U test was used to compare groups with non-normal distribution of quantitative values. For all statistical tests, significance was defined as p≤0.05. Analyses were conducted using the rstatix package (version 0.7.2) in R.^35^ Given the exploratory nature of this study, we did not correct for multiple comparisons across different statistical tests used.

K means clustering followed by principal curve projection were used to calculate significance of cluster separation in alpha diversity analyses. Analyses were performed using the factoextra (version 1.0.7) and ClusterSignificance (version 1.36.0) packages in R. ^32,33^

Complete code is available at: https://github.com/mcclur51e/METRCanalysis

## RESULTS

Intraoperatively collected samples were submitted for routine pathology and V1-V3 region 16S rRNA gene amplicon sequencing (patient demographics, wound characteristics, and clinical outcomes are summarized in Tables 1 and 2). Microbial culture results provided genus-level identification of bacterial isolates. 16S rRNA gene amplicon sequencing results provided amplicon sequence variant (ASV) identification of bacterial communities from extracted DNA. The combined microbial culture and 16S rRNA gene sequencing data were used to assess the association between microbial bioburden, culture results, and clinical outcomes. Samples were additionally divided into two groups based on collection timepoint: baseline (definitive wound closure) and follow-up (corrective surgery).

### Relationship Between Bioburden and Culture Positivity

#### Percent Bacterial Reads

ASVs from NGS were identified as bacterial or animal. Percent bacterial reads was calculated as the fraction of total NGS reads identified as bacterial. Samples that were culture-positive had a significantly higher median percent bacterial reads compared to culture-negative samples (62.8% ± 33.8% vs. 4.5% ± 21.4%, Wilcoxon rank-sum *p* = 1.6e-6). This difference remained statistically significant when analyzed by timepoint: baseline (*p* = 2e-4) and follow-up (*p* = 1.6e-3) (Figure 1A). A threshold value calculated to optimize Youden’s J statistic resulted in 67% sensitivity and 91% specificity of percent bacterial reads for predicting culture positivity (Youden = 0.58). These results suggest a strong positive association between NGS-derived bacterial read percentage and conventional culture results.

**Fig. 1.**
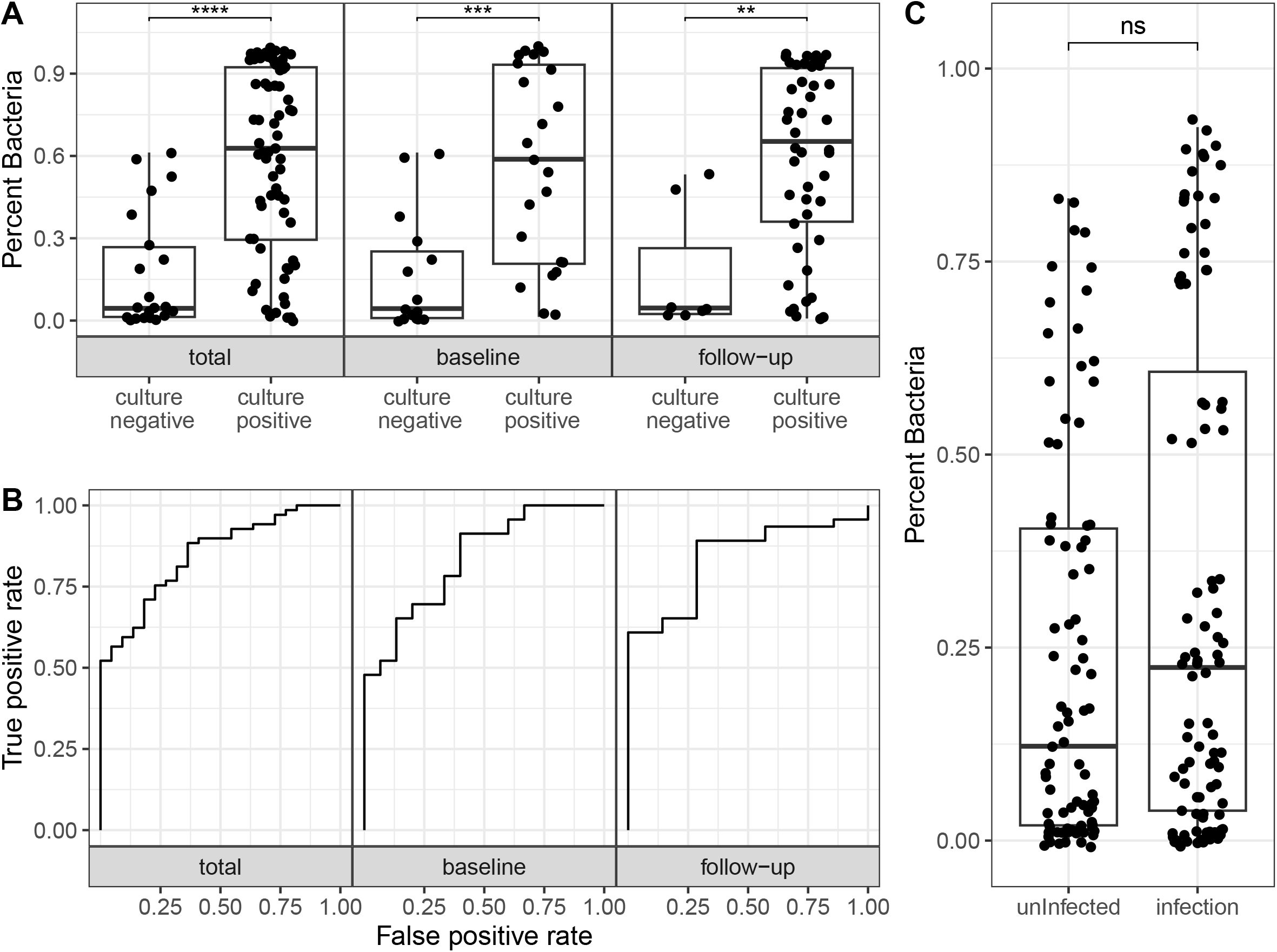
Percent of ASVs of intrasurgical tissue samples that are assigned to bacteria. A) Samples grouped by collection timepoint and qualitative culture result. B) Receiver operating curve for prediction of culture result based on percent bacterial ASVs. C) Samples grouped by outcome. Values compared using a Mann-Whitney U test (^*^ p≤0.05, ^**^ p≤0.01, ^***^ p≤0.001).

#### Bacteria-to-Human Reads Ratio

NGS data was further used to calculate the ratio of bacterial to human reads. The log of this bacterial to human ratio, log(BHR), was then calculated to normalize the data. Culture-positive samples exhibited a significantly higher log(BHR) than culture-negative samples (0.02 ± 0.94 vs.-1.55 ± 0.82, *p* = 1e-6). This difference was statistically significant at both timepoints (baseline *p* = 4e-4, follow-up *p* = 1e-3) (Figure 2A). A threshold value calculated to optimize Youden’s J statistic resulted in a 68% sensitivity and 91% specificity of log(BHR) for predicting culture positivity (Youden = 0.59). These findings support the utility of the log(BHR) as a semi-quantitative marker of microbial bioburden.

**Fig. 2.**
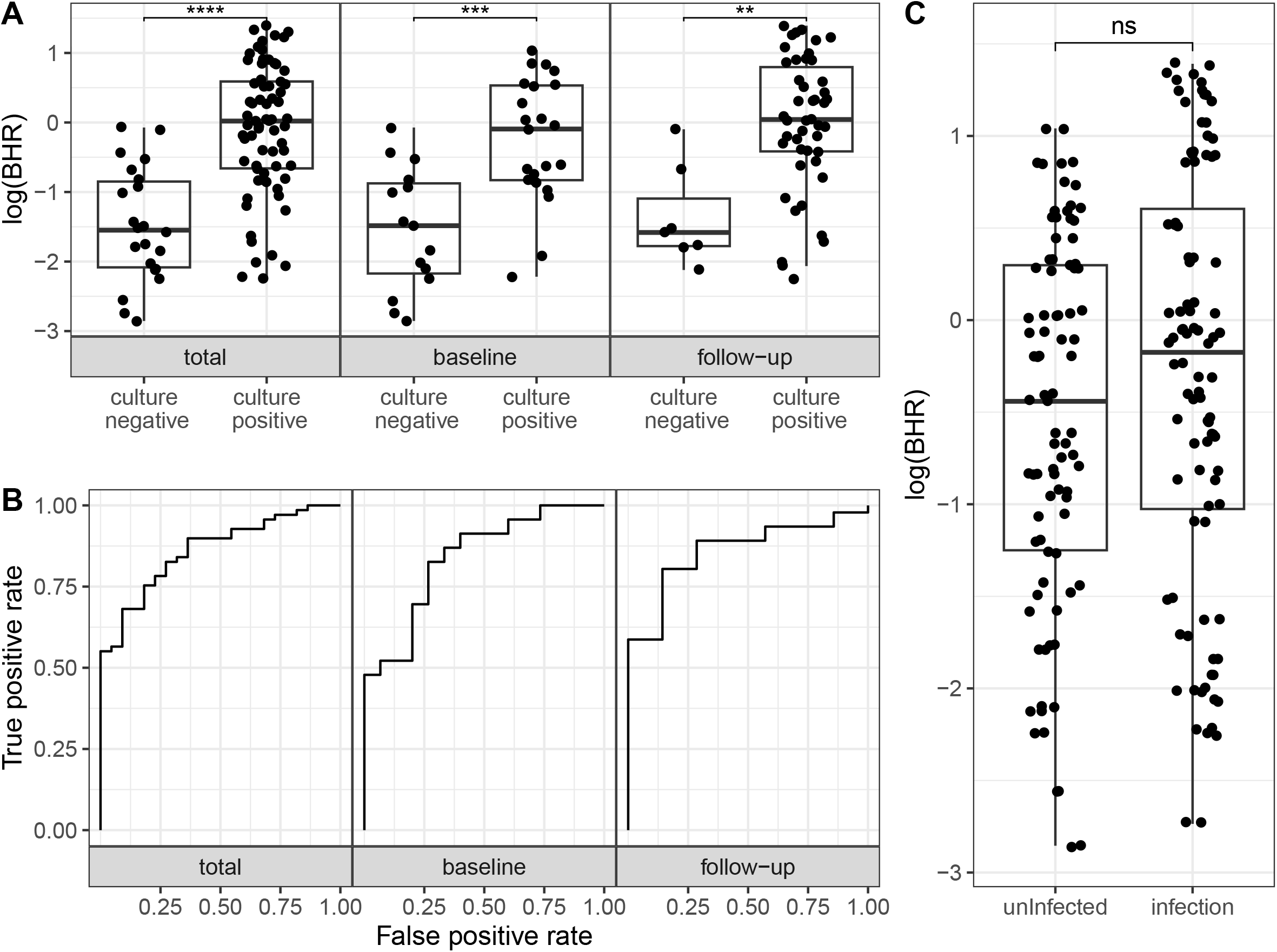
Bacteria-to-Human reads ratio (BHR) from 16S rRNA gene amplicon sequencing of intrasurgical tissue samples. A) Samples grouped by collection timepoint and qualitative culture result. B) Receiver operating curve for prediction of culture result based on BHR. C) Samples grouped by outcome. Note that y-axis is in log scale. Values compared using a Mann-Whitney U test (^*^ p≤0.05, ^**^ p≤0.01, ^***^ p≤0.001).

### Relationship Between Bioburden and Infection Incidence

#### Culture and NGS Positivity at Baseline

Neither baseline culture positivity nor baseline 16S rRNA gene amplicon sequencing positivity were associated with the later development of infection. 8 of 23 (34.8%) patients with positive baseline cultures and 7 of 15 patients (46.7%) with negative cultures (Table 1) developed infection. Similarly, 6 of 23 (26.1%) patients with NGS-positive baseline samples developed infection, compared to 9 of 15 (60.0%) with NGS-negative samples. The relationship between culture and NGS-positivity at baseline is significant (McNemar’s chi-squared test, p = 0.043).

#### Culture and NGS Positivity at Follow-Up

Follow-up culture positivity (often an infection diagnostic criterion) and 16S rRNA gene amplicon sequencing positivity increased in infection samples. Of the 46 patients with positive follow-up cultures, 32 (9.6%) were diagnosed with an infection and only 1 of 7 patients with negative cultures (14.3%). 31 of 49 (63.3%) patients with NGS-positive samples and 2 of 4 (50.0%) patients with NGS-negative samples at follow-up developed infection (Table 1). When results from culture and NGS are examined concomitantly, 30 of 44 (68.2%) patients with dual-positive samples at follow-up were diagnosed with infection and 0 of 2 (0%) of dual-negative patients. The relationship between culture and NGS-positivity at follow-up is significant (McNemar’s chi-squared test, p = 0.009). These results suggest that bioburden markers are more predictive in established infections than at baseline.

#### Quantitative Bioburden Measures

Samples from patients who developed infection exhibited no difference in bacterial read percentage (infection 58.7% SD 32.5%, no infection 43.4% SD 25.9%), *p* = 0.233, Figure 1C) and log(BHR) (infection −0.44 SD 1.01, no infection −0.18 SD 1.14), *p* = 0.117, Figure 2C). When examining timepoints individually, samples at baseline (p=0.281 and p=0.156 respectively) continued to exhibit no difference in bacterial read percentage or logBHR while samples at follow-up did show significant differences (p=0.006 and p=0.012 respectively) (Table S3).

### Alpha Diversity and Infection

Shannon and Simpson alpha diversity metrics are commonly used in microbiome research to quantify the taxonomic diversity within a sample. Shannon diversity accounts for the number of different species and evenness while Simpson diversity emphasizes how much one or a few species dominate the community. We sought to understand if microbial alpha diversity associated with clinical and culture features of our cohort.

#### Alpha Diversity

Both alpha diversity metrics at baseline showed no differences between patient outcomes (Figure 3). At follow-up, we found that our data indicated threshold values for both Shannon and Simpson diversity metrics above which all samples were from patients diagnosed with an infection (Figure 3A & B). High log(BHR), high Shannon diversity, and high Simpson diversity were each individually 100% specific for infection but had low sensitivity (38.7%, 32.3%, and 25.8%, respectively). Use of all three metrics in combination improved sensitivity to 58.1%. An additional small cluster was identified containing six infection samples with low log(BHR) and low Shannon or Simpson diversities (specificity=85.7%, sensitivity=9.6%). In combination, the two clusters resulted in a total sensitivity of 67.7%.

**Fig. 3.**
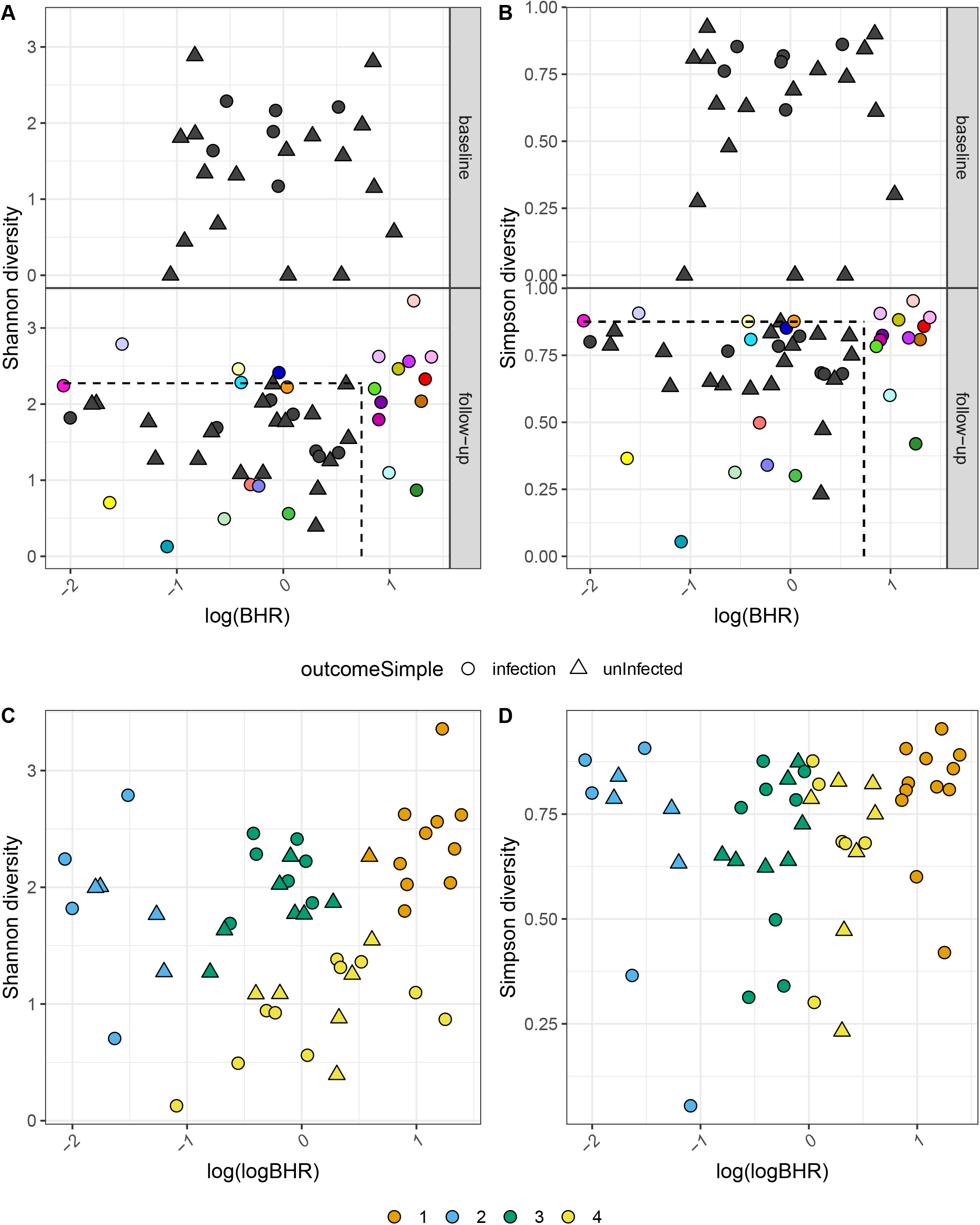
Plots showing A) Shannon and B) Simpson diversities as a function of Bacteria-to-Human reads ratio, log(BHR), in intrasurgical tissue samples. Plots are split by timepoint of sample collection. Colored points indicate the same sample between plots and highlights infections predicted by at least one of the three plotted metrics. Dashed lines indicate regions corresponding to clusters discussed in the results. Unsupervised cluster analyses of the same plots for follow-up samples are in C) Shannon and D) Simpson with colors indicating assigned clusters 1-4 (numbers are arbitrary). Cluster 1 in both plots (orange) includes many of the same infection samples highlighted in A and B. Projected separation p-value≤0.01 for all clusters. circles = infection, triangles=uninfected.

#### Machine Learning for Cluster Analysis

We further used K means clustering and a principal curve projection (an unsupervised learning algorithm) to partition follow-up samples into clusters based on alpha diversity. The results were similar for both Shannon and Simpson diversity metrics when plotted against log(BHR). One cluster consisted of 90.0-100% infection samples (9-12/12) with remaining clusters consisting of 44-63% infection samples (Figure 3C & D). This cluster corresponded to all 12 infection samples identified using high threshold log(BHR) values. No cluster corresponded to the infection samples identified using low threshold alpha diversity values. Separation of all four clusters was significant with p-value≤0.01. These results suggest that half of established infections are likely to have high bioburden, high dominance, and/or high evenness of their microbial community. With this small dataset we also were able to identify a pattern of low bioburden, low dominance, and/or low evenness of the microbiome in a small subset of established infections. However, this pattern was not statistically significant and requires a larger dataset to confirm.

## DISCUSSION

This study demonstrates that culture positivity in open fracture wounds is associated with increased overall bioburden, as quantified by next-generation sequencing (NGS)-based metrics. In addition to confirming the relationship between culture positivity and bioburden, our analysis identified two distinct microbial profiles in established infections. One cluster demonstrated high bioburden with high alpha diversity, consistent with polymicrobial infection and abundant bacterial cells. The second (smaller) cluster exhibited low alpha diversity and lower bioburden, suggestive of dominance by a single pathogen that either establishes infection with less bacterial cells or may by partially suppressed in expansion by treatment or the immune response (Figure 3). These disparate profiles may reflect different types of infection and could have implications for treatment strategies. Notably, baseline culture results and quantitative bioburden measures were not significantly associated with subsequent infection in this cohort, likely due to the limited sample size and reduced statistical power in this study. However, our findings lay the groundwork for future studies that are fully powered to validate predictive bioburden thresholds and microbial features that signal risk for infection.

Prior studies have shown that culture positivity at the time of definitive wound closure is associated with increased risk of subsequent infection. However, a persistent challenge has been the frequent mismatch between the pathogens identified in the open fracture and those isolated at time of later infection.^15,17,18^ This discordance has led to uncertainty in interpreting the clinical relevance of early open fracture culture results. One hypothesis is that culture positivity may serve, not as a predictor of specific pathogens, but rather as a surrogate marker for overall wound bioburden. Some have even proposed using cultures to guide the need for repeat debridement.^19^ However, this approach has not been widely adopted, in part due to ambiguity around what positive culture actually indicates.

Our findings provide molecular evidence supporting the interpretation that culture positivity reflects increased total bacterial load, rather than established infection. While this study was not powered to detect a statistically significant association between either culture or bioburden and subsequent infection, the observed trends combined with prior evidence linking culture positivity to infection suggest that such relationships may become clearer in larger studies. Importantly, our results underscore the potential for quantitative, sequencing-based bioburden assessment to offer more objective and actionable guidance than traditional culture. In contrast to culture, 16S rRNA gene amplicon sequencing-derived assessment may allow for the development of bioburden thresholds to inform more precise clinical decisions about timing of wound closure, need for further debridement, or use of local antimicrobials, without the interpretive confusion surrounding infection status.

The identification of two distinct microbial profiles—one characterized by high alpha diversity and high bioburden (suggestive of polymicrobial infections with a high burden of pathogens), and the other by lower diversity and lower bioburden (suggestive of dominance by a single pathogen with a smaller quantity of pathogens)—is a novel and potentially important finding. Clinically, this suggests that infections may follow distinct ecological trajectories, with implications for diagnosis, management, and research design. First, it reinforces the value of NGS not just for species identification, but for characterizing broader features of microbial ecology, such as diversity and community structure. While much of the current clinical enthusiasm around NGS has focused on using it to identify pathogens not captured by culture, our findings suggest that incorporating metrics like bioburden and alpha diversity may provide additional diagnostic and prognostic value. Second, these two patterns may reflect fundamentally different biological processes, such as early vs. established infection, or host-impaired immunity vs. pathogen virulence, and may respond differently to surgical and antimicrobial treatment. This highlights the need to stratify patients by microbial profile when evaluating treatment outcomes, as pooling them may obscure clinically meaningful differences in efficacy. Third, this approach may also inform risk prediction and tailored therapy. For example, high-diversity/high-burden infections may be more resistant to treatment and require broader-spectrum or prolonged antimicrobial strategies, while low-diversity infections dominated by a single pathogen might benefit from targeted therapy. Finally, the observation that alpha diversity and bioburden together improved specificity and sensitivity for infection suggests these metrics could ultimately contribute to a more nuanced, multi-parametric diagnostic framework, potentially guiding decisions around wound closure timing, need for re-debridement, or escalation of care.

In this context, our study quantitatively links culture positivity in open fracture wounds to increased total microbial bioburden using NGS-derived metrics. This molecular evidence supports the long-held—but previously unproven—hypothesis that early culture positivity reflects a higher bacterial burden, rather than the presence of specific infecting pathogens. It also highlights the potential utility of integrating NGS-based approaches into clinical workflows, not only for species identification but for real-time assessment of bioburden, diversity, and ecological shifts that may inform prognosis and treatment.

Our findings are consistent with and build upon a growing body of literature using molecular techniques to characterize microbial dynamics in open fracture wounds. Prior studies have shown that microbial communities are not static but evolve over time in response to both injury and treatment.^29–31^ A prospective study of 30 patients with open fracture used high throughput 16S rRNA gene amplicon sequencing to examine bacterial communities in open fractures longitudinally, demonstrating that microbiota composition became increasingly similar to adjacent skin microbiota with healing. This microbial convergence appeared to be shaped by injury mechanism, severity, anatomical site, and the occurrence of infectious complications. These findings suggest that clinical variables significantly influence microbial dynamics and that microbial signatures may eventually serve as biomarkers of healing trajectory or complication risk.^29^ A larger prospective cohort study of 155 patients with open fracture similarly used 16S rRNA gene amplicon sequencing to demonstrate that microbiota diversity decreased after infection onset, with a shift toward dominant pathogenic organisms like *Enterobacter* and *Pseudomonas*. Pre-infection open fracture samples in this study included *Bacillus* and *Staphylococcus* species, supporting the concept that specific early microbial patterns may signal susceptibility to future infection while not specifically identifying future pathogenic community members.^30^ Additionally, work from a separate prospective cohort of 52 open fracture patients further supports the dynamic nature of wound microbiota. In that study, microbial profiles from wounds and adjacent skin converged over time, and mechanism of injury emerged as the most important factor distinguishing microbial communities during healing. These findings align with our hypothesis that the open fracture microbiome is shaped not only by exogenous contamination but also by host and injury factors—offering a more nuanced view of infection pathogenesis than is afforded by culture-based methods alone.^31^ Together, these studies, including ours, point toward a future in which microbial profiling may offer actionable insights into wound management. With further validation in larger cohorts, it may be possible to define microbial thresholds or patterns that predict infection risk and guide the timing of wound closure, need for further debridement, or targeted antimicrobial strategies.

There are several strengths of this study. First, it leveraged a high-quality, prospectively collected dataset from a large, multicenter clinical trial with standardized data collection methods and rigorous outcome adjudication. This design minimizes recall and reporting bias and ensures consistent documentation of clinical variables and infectious complications. Second, the multicenter nature of the study, drawing from diverse Level I trauma centers across North America, enhances the generalizability of the findings by capturing variation in patient demographics, injury patterns, clinical practices, and geographic microbial exposures. This diversity allows for broader applicability of the conclusions beyond a single institution or region. Finally, the identification of distinct microbial signatures associated with infection offers a foundation for future research aimed at developing targeted diagnostics and personalized treatment strategies.

There are several limitations associated with this study. First, this is a secondary analysis of data collected as part of a larger prospective cohort study that was not specifically designed to investigate NGS-based microbiome features or bioburden. As such, while the dataset includes high-quality clinical data, the timing and context of sample collection may not be optimized for mechanistic microbiome analysis, and confounding is possible. Second, while the use of NGS offers improved resolution over traditional culture, there are inherent limitations to NGS-based approaches. These include potential contamination during DNA extraction or amplification, variable sequencing depth across samples, biases of sequencing primers, and the inability to distinguish between viable and non-viable organisms. Additionally, while we used bacterial-to-human reads ratios and bacterial read percentages as proxies for bioburden, these are indirect measures and may be influenced by factors such as tissue type, sample biomass, or host response variability. Third, the relatively small sample size, particularly in the subgroup that developed infection, limits the power to detect significant associations between baseline bioburden and clinical outcomes, and increases the risk of type II error. Fourth, although the prospective, multi-center nature of the parent trial supports generalizability, all sites were Level I or II trauma centers with robust infrastructure, which may limit external validity to less-resourced settings. Taken together, these limitations underscore the need for future prospective studies with dedicated sample collection protocols, larger and more diverse cohorts, and standardized sequencing and analytic pipelines to validate and expand upon these findings.

In conclusion, this study provides evidence that sequencing-based bioburden metrics correspond closely with traditional culture positivity in open fractures. By quantifying total microbial load and identifying distinct infection profiles, NGS-derived measures may move beyond the limitations of culture to offer a more nuanced understanding of wound microbiology. These findings support the hypothesis that bioburden, not necessarily the presence of specific pathogens, is a critical determinant of infection risk. As such, the integration of quantitative microbial profiling into clinical workflows has the potential to improve timing and precision of surgical decision-making. Future prospective studies in larger cohorts are essential to validate bioburden thresholds and microbial community patterns that can be used to guide personalized treatment strategies and ultimately reduce the persistently high rate of infectious complications in patients with open fractures.

## Supporting information

Corporate Author List

## ACKNOWLEDGEMENTS

The authors wish to thank members of the Ross lab for providing feedback on the manuscript. This work is supported by National Institutes of Health grant R00GM129874 and R35GM142685 to BDR and Department of Defense Orthopedic Extremity Trauma Research Program (DOD OETRP) W81XWH-09-20108 to METRC.

## Author contributions

METRC collected, cultured, and sequenced all samples. EAM performed all data analysis. EAM, LG, and BDR wrote the manuscript. All authors reviewed and edited the manuscript.

## Data and material availability

Code used for processing sequencing data is available at https://github.com/mcclur51e/METRCanalysis

## TABLES

**Table S1.** Patient demographics.

**Table S2.** Wound characteristics.

**Table S3.** 16S rRNA gene amplicon sequencing-determined bioburden of intrasurgical tissue samples. Medians and standard deviations are calculated by timepoint and final patient diagnosis. Mann-Whitney U test p-values comparing groups in each row (all p-values are not significant).

## REFERENCES

1. Bosse MJ, MacKenzie EJ, Kellam JF, et al. An analysis of outcomes of reconstruction or amputation after leg-threatening injuries. N Engl J Med. 2002;347(24):1924–1931. doi:10.1056/NEJMoa012604

2. Yun HC, Murray CK, Nelson KJ, Bosse MJ. Infection After Orthopaedic Trauma: Prevention and Treatment. J Orthop Trauma. 2016;30 Suppl 3:S21–S26. doi:10.1097/BOT.0000000000000667

3. Harris AM, Althausen PL, Kellam J, Bosse MJ, Castillo R, Lower Extremity Assessment Project (LEAP) Study Group. Complications following limb-threatening lower extremity trauma. J Orthop Trauma. 2009;23(1):1–6. doi:10.1097/BOT.0b013e31818e43dd

4. Keeling JJ, Gwinn DE, Tintle SM, Andersen RC, McGuigan FX. Short-term outcomes of severe open wartime tibial fractures treated with ring external fixation. J Bone Joint Surg Am. 2008;90(12):2643–2651. doi:10.2106/JBJS.G.01326

5. Lerner A, Fodor L, Soudry M. Is staged external fixation a valuable strategy for war injuries to the limbs? Clin Orthop Relat Res. 2006;448:217–224. doi:10.1097/01.blo.0000214411.60722.f8

6. Owens BD, Kragh JF, Wenke JC, Macaitis J, Wade CE, Holcomb JB. Combat wounds in operation Iraqi Freedom and operation Enduring Freedom. J Trauma. 2008;64(2):295–299. doi:10.1097/TA.0b013e318163b875

7. Merritt K. Factors increasing the risk of infection in patients with open fractures. J Trauma. 1988;28(6):823–827.

8. Dellinger EP, Miller SD, Wertz MJ, Grypma M, Droppert B, Anderson PA. Risk of infection after open fracture of the arm or leg. Arch Surg. 1988;123(11):1320–1327.

9. Premkumar A, Kolin DA, Farley KX, et al. Projected Economic Burden of Periprosthetic Joint Infection of the Hip and Knee in the United States. J Arthroplasty. 2021;36(5):1484–1489.e3. doi:10.1016/j.arth.2020.12.005

10. Olesen UK, Pedersen NJ, Eckardt H, et al. The cost of infection in severe open tibial fractures treated with a free flap. Int Orthop. 2017;41(5):1049–1055. doi:10.1007/s00264-016-3337-6

11. Morcos MW, Kooner P, Marsh J, Howard J, Lanting B, Vasarhelyi E. The economic impact of periprosthetic infection in total knee arthroplasty. Can J Surg. 2021;64(2):E144–E148. doi:10.1503/cjs.012519

12. Hackett DJ, Rothenberg AC, Chen AF, et al. The economic significance of orthopaedic infections. J Am Acad Orthop Surg. 2015;23 Suppl:S1–7. doi:10.5435/JAAOS-D-14-00394

13. Barnes CL. Overview: the health care burden and financial costs of surgical site infections. Am J Orthop (Belle Mead NJ). 2011;40(12 Suppl):2–5.

14. Trampuz A, Zimmerli W. Diagnosis and treatment of infections associated with fracture-fixation devices. Injury. 2006;37 Suppl 2:S59–66. doi:10.1016/j.injury.2006.04.010

15. Burns TC, Stinner DJ, Mack AW, et al. Microbiology and injury characteristics in severe open tibia fractures from combat. Journal of Trauma and Acute Care Surgery. 2012;72(4):1062–1067. doi:10.1097/TA.0b013e318241f534

16. Lingaraj R, Santoshi JA, Devi S, et al. Predebridement wound culture in open fractures does not predict postoperative wound infection: A pilot study. J Nat Sci Biol Med. 2015;6(Suppl 1):S63–68. doi:10.4103/0976-9668.166088

17. Lee J. Efficacy of Cultures in the Management of Open Fractures. Clinical Orthopaedics and Related Research (1976-2007). 1997;339:71–75.

18. Valenziano CP, Chattar-Cora D, O’Neill A, Hubli EH, Cudjoe EA. Efficacy of primary wound cultures in long bone open extremity fractures: are they of any value? Arch Orthop Trauma Surg. 2002;122(5):259–261. doi:10.1007/s00402-001-0363-6

19. Lenarz CJ, Watson JT, Moed BR, Israel H, Mullen JD, Macdonald JB. Timing of wound closure in open fractures based on cultures obtained after debridement. J Bone Joint Surg Am. 2010;92(10):1921–1926. doi:10.2106/JBJS.I.00547

20. Major Extremity Trauma Research Consortium (METRC). The Bioburden Associated with Severe Open Tibial Fracture Wounds at the Time of Definitive Closure or Coverage: The BIOBURDEN Study. J Bone Joint Surg Am. Published online March 15, 2024. doi:10.2106/JBJS.23.00157

21. Stennett CA, O’Hara NN, Sprague S, et al. Effect of Extended Prophylactic Antibiotic Duration in the Treatment of Open Fracture Wounds Differs by Level of Contamination. J Orthop Trauma. 2020;34(3):113–120. doi:10.1097/BOT.0000000000001715

22. Murray C. Infectious disease complications of combat-related injuries. Critical care medicine. Published online July 2008. doi:10.1097/CCM.0b013e31817e2ffc

23. Okike K, Bhattacharyya T. Trends in the management of open fractures. A critical analysis. J Bone Joint Surg Am. 2006;88(12):2739–2748. doi:10.2106/JBJS.F.00146

24. Mangram AJ, Horan TC, Pearson ML, Silver LC, Jarvis WR. Guideline for Prevention of Surgical Site Infection, 1999. Centers for Disease Control and Prevention (CDC) Hospital Infection Control Practices Advisory Committee. Am J Infect Control. 1999;27(2):97–132; quiz 133-134; discussion 96.

25. McClure EA, Nelson MC, Lin A, Graf J. Macrobdella decora: Old World Leech Gut Microbial Community Structure Conserved in a New World Leech. Appl Environ Microbiol. 2021;87(10):e02082–20. doi:10.1128/AEM.02082-20

26. Callahan B, McMurdie P, Rosen M, Han A, Johnson A, Holmes S. DADA2: High-resolution sample inference from Illumina amplicon data. Nature methods. Published online July 2016. doi:10.1038/nmeth.3869

27. Pruesse E, Peplies J, Glöckner FO. SINA: accurate high-throughput multiple sequence alignment of ribosomal RNA genes. Bioinformatics. 2012;28(14):1823–1829. doi:10.1093/bioinformatics/bts252

28. Walker SP, Barrett M, Hogan G, Flores Bueso Y, Claesson MJ, Tangney M. Non-specific amplification of human DNA is a major challenge for 16S rRNA gene sequence analysis. Sci Rep. 2020;10(1):16356. doi:10.1038/s41598-020-73403-7

29. Hannigan GD, Grice EA. Microbial ecology of the skin in the era of metagenomics and molecular microbiology. Cold Spring Harb Perspect Med. 2013;3(12):a015362. doi:10.1101/cshperspect.a015362

30. Sukpanichyingyong S, Sae-Jung S, Stubbs DA, Luengpailin S. Microbiota shifts in fracture-related infections and pathogenic transitions identified by 16S rDNA sequencing. Sci Rep. 2025;15(1):7732. doi:10.1038/s41598-025-91990-1

31. Bartow-McKenney C, Hannigan GD, Horwinski J, et al. The microbiota of traumatic, open fracture wounds is associated with mechanism of injury. Wound Repair and Regeneration. 2018;26(2):127–135.

32. Kassambara A, Mundt F (2020). factoextra: Extract and Visualize the Results of Multivariate Data Analyses. doi:10.32614/CRAN.package.factoextra < https://doi.org/10.32614/CRAN.package.factoextra >, R package version 1.0.7, < https://CRAN.R-project.org/package=factoextra >.

33. Serviss JT, Gadin JR, ERiksson P, Folkersen L, Grander D. ClusterSignificance: a bioconductor package facilitating statistical analysis of class cluster separations in dimensionality reduced data. Bioinformatics. 2017;33(19): 3126–3128.

34. Thiele C, Hirschfeld G. cutpointr: Improved estimation and validation of optimal cutpoints in R. Journal of Statistical Software. 2021. 98(11).

35. Kassambara A (2023). rstatix: Pipe-Friendly Framework for Basic Statistical Tests. R package version 0.7.2, https://rpkgs.datanovia.com/rstatix/.

