## Supplementary material for "Culture Positivity Reflects Quantitative Bacterial Bioburden in Open Fractures": Corporate Author List

Emily Ann McClure^1^, Roman M. Natoli, MD, PhD^2^, Renan C. Castillo, PhD^3^,
Anthony R. Carlini, MS^3^, Joseph C. Wenke, PhD^4^, Robert V. O'Toole, MD^5^,
Gregory M. Schrank, MD, MPH^6^, William Obremskey, MD, MPH^7^,
Stephen J. Warner, MD, PhD^8^, Michael J. Bosse, MD^9^, Benjamin D. Ross, PhD,^1,10*^,
Ida Leah Gitajn, MD, MS^10*^and METRC^3^

**Institutional Affiliations:**

^1^Department of Microbiology and Immunology, Geisel School of Medicine, Dartmouth College, Hanover, NH

^2^Department of Orthopedic Surgery, Indiana University Health Methodist Hospital, Indianapolis, IN

^3^Department of Health Policy and Management, Johns Hopkins Bloomberg School of Public Health, Baltimore, MD

^4^Department of Orthopaedic Surgery and Rehabilitation, University of Texas Medical Branch & Shriners Children's Texas, Galveston, TX

^5^Department of Orthopaedics, University of Maryland School of Medicine, Baltimore, MD

^6^Department of Epidemiology & Public Health, University of Maryland School of Medicine Baltimore, MD

^7^Department of Orthopaedic Surgery, Vanderbilt University Medical Center, Nashville, TN

^8^Department of Orthopedic Surgery, McGovern Medical School at The University of Texas Health Science Center at Houston, Houston, TX

^9^Department of Orthopaedic Surgery, Atrium Health Carolinas Medical Center, Charlotte, NC

^10^Department of Orthopedics, Geisel School of Medicine, Dartmouth College, Hanover, NH

**Corresponding Author**:

First Last, MD

123 Main St, City, ST 12345

**Conflicts of Interest and Source of Funding**:

The authors have no actual or potential conflicts of interest to disclose. Work on the original BIOBURDEN study was supported by the United States Department of Defense Grant # W81XWH-09-20108. The original sponsor had no role in the design or conduct of the trial; the collection, management, analysis, or interpretation of the data; or the preparation, review, or approval of this manuscript for submission.

**METRC Corporate Author Appendix:
*The following individuals meet ICMJE criteria for authorship. Site affiliations are as of time of study unless otherwise noted****.*

*Atrium Health-Carolinas Medical Center*: Joseph R. Hsu, MD; Madhav A. Karunakar, MD; Rachel B. Seymour, PhD; Stephen H. Sims, MD; Christine Churchill, MA; *Atrium Health Wake Forest Baptist*: Eben A. Carroll, MD; Martha B. Holden, AAS, AA; *Barnes-Jewish Hospital at Washington University in St. Louis*: Anna N. Miller, MD (now affiliated with Dartmouth Hitchcock Clinic); Amanda Hughes LeFranc, PhD (now affiliated with UConn Health); *Boston Medical Center*: Paul Tornetta III, MD, PhD; *Brooke Army Medical Center*: Daniel J. Stinner, MD, PhD (now affiliated with Vanderbilt University Medical Center); Clinton K. Murray, MD; Heather C. Yun, MD (now affiliated with South Texas Veterans Healthcare System); *Center for Genomic Sciences, Allegheny Singer Research Institute*: Garth D. Ehrlich	PhD, FAAAS (no longer affiliated); *Center for Orthopaedic Research and Education*: Clifford B. Jones, MD, FACS (now affiliated with Creighton University School of Medicine Phoenix); Debra L. Sietsema, MSN, PhD (no longer affiliated); *Denver Health and Hospital Authority*: Corey Henderson Trujillo, DO (now affiliated with Legacy Health); *Duke University Medical Center*: Rachel M. Reilly, MD; Robert D. Zura, MD (now affiliated with Louisiana State University); Cameron Howes, BA; *Florida Orthopaedic Institute at Tampa General Hospital*: Hassan Mir, MD, MBA; *Grant Medical Center*: Benjamin C. Taylor, MD, PhD; *Hennepin Healthcare*: Patrick Yoon, MD (now affiliated with University of Minnesota); *Indiana University Health Methodist Hospital*: Anthony Sorkin, MD (now affiliated with Indiana Orthopedic Institute); Walter W. Virkus, MD; *Louisiana State University*: Olivia C. Lee, MD; *McGovern Medical School at The University of Texas Health Science Center at Houston*: Joshua L. Gary, MD (now affiliated with Keck School of Medicine of USC); Mark R. Brinker, MD; Andrew Choo, MD; John W. Munz, MD; Matthew C. Galpin, CCRC (now affiliated with Profits Prophet); *MetroHealth Medical Center*: Heather A. Vallier, MD (now affiliated with Cleveland Clinic Foundation) *Mission Hospital*: H. Michael Frisch, MD (now affiliated with Novant Health Charlotte Orthopedic Hospital); Thomas M. Large, MD (now affiliated with Emory University School of Medicine); *Naval Medical Center Portsmouth*: Christopher S. Smith, MD, MBA; *Orlando Regional Medical Center*: Joshua Langford, MD; *R Adams Cowley Shock Trauma Center at the University of Maryland*: Theodore Manson, MD, MS; W. Andrew Eglseder, MD (now Emeritus); *Rhode Island Hospital at Brown University*: Roman Hayda, MD; *Ryder Trauma Center at the University of Miami*: Gregory A. Zych, DO; *Stanford University Medical Center*: Michael J. Gardner, MD; *Temple University Hospital*: Saqib Rehman, MD, MBA; *The University of California, San Francisco*: Theodore Miclau, MD; Eleni Berhaneselase, BA; *University of Mississippi Medical Center*: Patrick F. Bergin, MD; Clay A. Spitler, MD (now affiliated with University of Alabama at Birmingham); *University of Oklahoma Medical Center*: David Teague, MD; *University of Virginia Health*: David B. Weiss, MD; *University of Washington Medicine Harborview Medical Center*: Reza Firoozabadi, MD; *Vanderbilt University Medical Center*: Robert Boyce, MD; Eduardo J. Burgos, MD (now affiliated with Fundación Santa Fe de Bogota); Andres Rodriguez-Buitrago, MD (now affiliated with Fundación Santa Fe de Bogota); Rajesh R. Tummuru, MBBS, MBA (no longer affiliated); Karen Trochez, MMHC; *Walter Reed National Military Medical Center*: Wade Gordon, MD (now affiliated with UNLV School of Medicine); Xochitl Ceniceros, PhD, CCRP; Sandra L. Waggoner, BA (now affiliated with Uniformed Services University); *METRC Coordinating Center at the Johns Hopkins Bloomberg School of Public Health*: Susan C. Collins, MSc

**Acknowledgements**:

***These individuals do not meet the ICMJE criteria for authorship, but the authors would like to outline their specific contributions to the project.***

The authors thank the following for their work on the original study: Christopher M. McAndrew, MD, MSc, Barnes-Jewish Hospital at Washington University Medical Center, for patient enrollment; Hope Carlisle, RN, BSN, Boston Medical Center (no longer affiliated), for patient enrollment; Mary Zadnik, ScD, MEd, OT, METRC Coordinating Center at the Johns Hopkins Bloomberg School of Public Health (now affiliated with St. Edwards University), for study design, data collection tools, team training, and site data collection monitoring; Amanda Amundson, MPH, MSc, R Adams Cowley Shock Trauma Center at the University of Maryland (no longer affiliated), for data acquisition; Lisa K. Cannada, MD, St. Louis University Hospital (now affiliated with University of North Carolina School of Medicine), for patient enrollment; Saam Morshed, MD, PhD, The University of California, San Francisco, for patient enrollment; John T. Gorczyca, MD, University of Rochester Medical Center, for patient enrollment and protocol review; Veronica Lester-Ballard, MSEd, MSN, University of Virginia Medical Center (no longer affiliated), for patient enrollment and follow up, and data acquisition; and Julie Agel, MA, University of Washington Medicine Harborview Medical Center (now affiliated with University of Minnesota), for data collection.
